# Machine learning identifies the immunological signature of Juvenile Idiopathic Arthritis

**DOI:** 10.1101/382499

**Authors:** Erika Van Nieuwenhove, Vasiliki Lagou, Lien Van Eyck, James Dooley, Ulrich Bodenhofer, An Goris, Stephanie Humblet-Baron, Carine Wouters, Adrian Liston

## Abstract

Juvenile idiopathic arthritis (JIA) is the most common childhood rheumatic disease, with a strongly debated pathophysiological origin. Both adaptive and innate immune processes have been proposed as primary drivers, which may account for the observed clinical heterogeneity, but few high-depth studies have been performed. Here we profiled the adaptive immune system of 85 JIA patients and 43 age-matched controls, identifying immunological changes unique to JIA and others common across a broad spectrum of childhood inflammatory diseases. The JIA immune signature was shared between clinically distinct subsets, but was accentuated in the systemic JIA patients and those patients with active disease. Despite the extensive overlap in the immunological spectrum exhibited by healthy children and JIA patients, machine learning analysis of the dataset proved capable of diagnosis of JIA patients with ~90% accuracy. These results pave the way for large-scale longitudinal studies of JIA, where machine learning could be used to predict immune signatures that correspond to treatment response group.

## Introduction

Juvenile idiopathic arthritis (JIA) is the most common childhood rheumatic disease. It is characterized by the onset of arthritis with no defined cause prior to 16 years of age, and the persistence of symptoms for more than six weeks. Evidence exists for roles of both genetic inheritance and the influence of unknown environmental triggers (1, 2). Elucidation of the precise etiology and pathogenesis of JIA remains complicated by the clinical heterogeneity in the group as a whole (3, 4). JIA patients were classified into 7 subtypes by the International League of Associations for Rheumatology (ILAR) according to clinical features, however it remains unknown as to whether each subtype has a distinct pathogenesis, and this classification may require revision with more complete understanding of the pathophysiology.

Detailed analysis of cellular immunophenotypes and genetic variants associated with JIA subtypes could help improve the current classification system (5). T cells are central to the pathogenesis of JIA and research has focused on unraveling the dynamic balance between pro-inflammatory (T helper 17 (Th17), Th1) and anti-inflammatory regulators (regulatory T cells (Treg)), but the debate on the driving effector CD4 helper subset remains ongoing (6-8). In particular, the inflammatory nature of IFNγ-producing Th1 cells in an arthritic context has been questioned, with IFNγ-deficient mice developing systemic JIA (sJIA)-like symptoms upon immune stimulation (9). High-depth immunophenotyping of the innate immune response to stimuli has recently identified a sJIA signature (10), however a similar in depth study on the adaptive immune response has been lacking. While most JIA subtypes share a strong similarity with autoimmune diseases, systemic JIA (sJIA) is often considered an autoinflammatory disease, raising the possibility of distinct immunological drivers of disease.

Identification of the immunological signature of JIA can substantially improve our understanding of the disease pathophysiology, can lead to better diagnosis and disease classification, and in the future may be used to stratify patients for appropriate therapeutic approaches. Here we performed deep immunophenotyping on a large cohort of unrelated JIA patients. We found JIA-specific and common inflammatory immune signatures, driving a distinct immunological profile to that of healthy children. The immunological signature of JIA was identifiable through machine learning and was elevated in systemic JIA and active JIA patients, leading to the potential future application of immune-led machine learning in JIA treatment selection trials.

## Methods

### Study population and sampling

Patients were recruited through the pediatric rheumatology department of Leuven University Hospital. All individuals or their legal guardians gave written informed consent, and the study was approved by the Ethics Committee of the University Hospitals Leuven. JIA patients were classified via three distinct systems. First, patients were classified based on the International League of Associations for Rheumatology (ILAR) classification criteria {Petty, 1998 #96} into persistent oligoarticular, extended oligoarticular, rheumatoid factor positive (RF+) polyarticular, rheumatoid factor negative (RF-) polyarticular, enthesitis-related, and systemic juvenile idiopathic arthritis (sJIA). Second, we used a classification system that combined polyarticular arthritis with extended oligoarticular arthritis, resulting in three categories: persistent oligoarticular, combined polyarticular / extended oligoarticular and systemic. Third, we used a classification system that first distinguished systemic patients, and then secondarily divided non-systemic patients on anti-nuclear antibody (ANA) status (using a cut-off titer of 160), resulting in three categories: ANA+ JIA, ANA-JIA and sJIA. Disease activity was assessed according to the Wallace criteria {Wallace, 2011 #97}. Disease controls consisted of two distinct populations, first, a group of patients diagnosed with juvenile-onset systemic lupus erythematosus, and second, a group of patients with non-arthritis systemic inflammatory diseases (including systemic autoinflammatory disorder and periodic fever autoinflammatory disease). The distribution of age at time of sampling in cases and controls is shown in **Supplementary Figure 1**. Blood samples from all participants were collected in heparin tubes and rested at 22 °C for 4 h before separation of serum and PBMCs using lymphocyte separation medium (LSM, MP Biomedicals). PBMCs were frozen in 10% DMSO (Sigma) and stored at -80 °C for a maximum of 10 weeks.

### Flow cytometry phenotyping

Thawed cells were stained with antibodies to allow deep immunophenotyping (**Supplementary Table 1**). Ki67 and FOXP3 staining was performed after treatment with fixation-permeabilization buffer (eBioscience). Cytokine staining was performed after *ex vivo* stimulation for 5 hours in 50ng/ml PMA (Sigma) and 500ng/ml ionomycin (Sigma) in the presence of GolgiStop (BD Biosciences). Stimulated cells were surface stained, fixed and permeabilized with Cytofix/Cytoperm (BD), before staining for cytokines. Data were acquired on a BD FACSCantoII and analyzed with FlowJo (Tree Star).

### Statistical analysis

Data on 42 immunological parameters (**Supplementary Table 2**), with a focus on cellular subsets within the adaptive immune system, were generated for 43 age-matched controls, 16 disease controls and 85 JIA patients (**see Supplementary Dataset 1 and 2**). Data (phenotypic, flow cytometric and serological) were collated and stored in Microsoft Excel. All data analysis was performed using R (http://www.r-project.org version 3.1.2 (11). The flow cytometry data were expressed as percentages as exported from FlowJo. Statistical comparison was based on Kruskal-Wallis one-way ANOVA followed by Dunn post-hoc test implemented in R and *P* values were adjusted with the False Discovery Rate method (**Supplementary Code 1 and 2**).

### Machine learning

In order to investigate the ability of machine learning to classify patient groups, the following model selection procedure was performed: the cohort was randomly split into ten almost equally large groups, so-called folds. Using these groups/folds, ten-fold cross validation was performed in order to assess each method’s ability to generalize to previously unseen cases. In every test, hyperparameter selection was performed on a training set consisting of nine folds. Then the best hyperparameters were used to train a model on the nine training folds, and the performance of this model was evaluated using the withheld test fold, which was neither used for training nor for the selection of hyperparameters. This procedure was performed for all ten folds. Dataset training was run using Random Forests, Artificial Neural Networks and Support Vector Machines, with the capacity of each to explain the data assessed on the basis of superior area under the ROC (receiver-operator characteristics).

Random Forests were run using hyperparameter selection with out-of-bag estimates on the respective training sets (12, 13). We considered 5 and 11 features per split. The sampling schemes trialed were: 1) Optimizing for accuracy, we considered the default sampling scheme which took the total number of samples, but the sampling was done with replacement. Moreover, sampling was performed independently of the samples’ class memberships. 2) Optimizing for balanced accuracy, the balanced random forest sampling scheme (14) was applied: each tree is based on as many positive samples as there are in the training set and exactly as many negative samples. 3) Use of choice of sampling scheme as a second hyperparameter, to select for superior area under the ROC curve: in addition to the schemes above, two more sampling schemes were added, where each tree is based on as many negative samples (smaller class) as there are in the training set and 1.5 and 2 times as many positive samples (larger class). Regardless of the goal criterion, we always constructed ensembles of 10,001 trees. Model selection was performed using R 3.3.0 (15) with the ‘randomForest’ package (16).

Artificial neural networks. Hyperparameter selection was performed using nine-fold cross validation on the respective training sets. The following hyperparameters were considered, with 672 combinations:

> Network architecture: we considered networks with one hidden layer consisting of 25, 50, 100 and 200 nodes as well as networks with two hidden layers with 25, 50, and 100 nodes each (with 7 different architectures in total).
>
> Number of training epochs: 150 and 300.
>
> Learning rates: 0.005 and 0.01.
>
> Momentum: 0.5 and 0.9.
>
> Dropout: dropout both for input nodes and hidden activations at rates of 0.2 and 0.5 (Srivastava et al., 2014).
>
> Classweights: the larger class was always weighted with 1 in the objective function, while we tried class weights of 1, 1.5, and 2 for the smaller class in order to allow the network to better deal with the unbalanced distribution of classes.
>
> Weight decay (aka L2 regularization): with and without weight decay with a regularization parameter of 0.001.

For all artificial neural networks we trained, we used ReLU activations (17) for the hidden nodes and a sigmoid activation for the output node along with binary cross-entropy as optimization objective. Training was performed by stochastic gradient descent using a mini-batch size of 32 samples. All computational experiments were implemented in Python 3 using the KERAS framework (18) as simple interface to the TensorFlow framework (19).

Potential Support Vector Machine was employed due to superior performance with unbalanced data (20). PSVM model selection was performed twice, once without balancing (to optimize for accuracy) and once with balancing (to optimize for balanced accuracy). Hyperparameter selection was performed using ten-fold cross validation on the respective training sets using all combinations of cost factors C ∈{8,10,12,…,20} and shrinkage parameters ε ∈{0.5,0.7,0.9,…,2.3}. In all experiments, Potential Support Vector Machine was used in dyadic mode, i.e. the model’s discriminant function is a linear combination of a subset of features (20).

## Results

### A distinct immunological signature of Juvenile Idiopathic Arthritis

To identify the potential immunological processes driving JIA, we recruited 85 JIA patients and 43 age-matched controls. JIA patients and healthy pediatric controls were assessed for 42 immunological parameters by flow cytometry, using the immune phenotyping platform previously validated (21). This platform gives a strong representation of key subsets of the adaptive immune system (**Supplementary Table 2**). As previously observed (21), immunological variance within the healthy control population was structured, with strong correlations between variance in different immunological parameters (**Figure 1A**). While much of this interrelatedness was preserved in JIA patients, for several adaptive immune subsets, such as iNKT cells, multiple disjunctions were observed (**Figure 1A**). For example, the homeostatic weak negative relationship between NKT cells and plasmablasts, observed in healthy controls, was inverted to a strong positive relationship in JIA, where increased frequencies of iNKT cells were associated with elevated plasmablasts (**Figure 1A**). Taking into account the variance in all assessed parameters, the immunological composition of JIA patients diverged from healthy controls, with overlapping but distinct separation of the two clusters on a PCA analysis (**Figure 1B**). To determine whether this JIA immune signature was a generic inflammatory disease signature, or a JIA-specific signature, we recruited 15 disease controls, juvenile patients with inflammatory disease but no arthritis, including 5 patients with juvenile-onset SLE and 10 patients with juvenile-onset systemic inflammatory diseases. PCA analysis of disease controls showed a little separation from the JIA clusters on a PCA plot (**Figure 1B**). As invariant traits compress PCA clusters, we reclustered individuals with a core immune set of 22 traits, limited to those showing significant variance (**Figure 1C**). In the core trait analysis, the JIA cluster showed separation from the disease control cluster, and both were in turn separated by a larger distance from the healthy cluster, albeit still with strong overlap between all three groups. These results demonstrate that juvenile inflammatory patients present with alterations in their immune profile, with both a core universal inflammatory signature and a JIA-specific immune signature underlying the separation.

**Figure 1.**
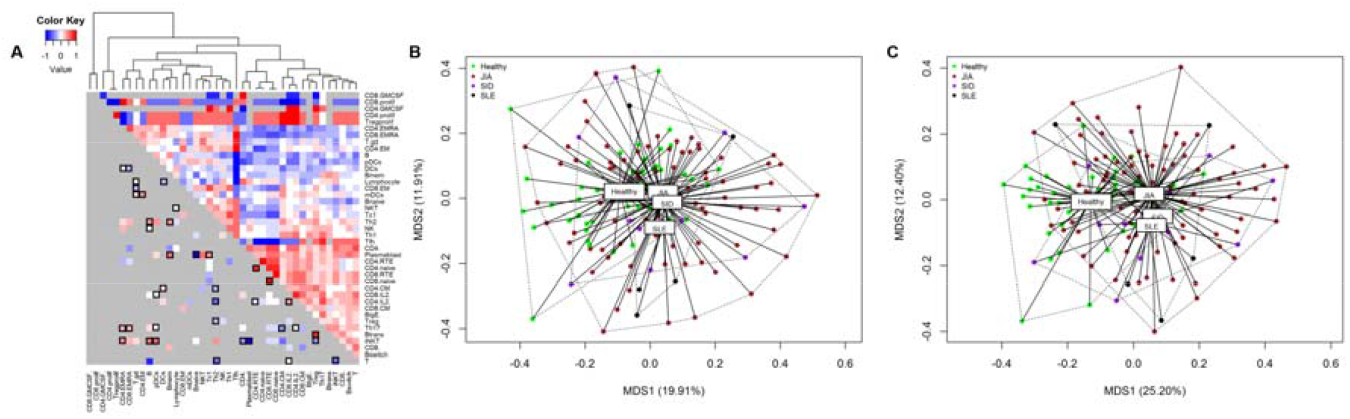
Juvenile Idiopathic Arthritis is marked by global immunological shifts and alteration in the relationship between leukocyte subsets. Healthy controls, JIA patients, SLE patients and systemic inflammatory disease (SID) patients were assessed for immune phenotype (n=43,86,16). (**A**) Upper right, above the diagonal: coregulation between pairs of cell types in healthy controls (n = 43) (red: positive correlation coefficient, blue: negative correlation coefficient, light gray: no data available). Unbiased clustering of coefficients was performed to group coregulated cell types. Lower left, below the diagonal: dark gray indicates coregulation between pairs of cell types in JIA patients (n=85) that are preserved from healthy controls. Coregulation between pairs of cell types that are significantly altered by disease (*p* < 0.05, and boxed if *p* < 0.01) is colored (red: positive correlation coefficient, blue: negative correlation coefficient). (**B**) All individuals were plotted with multidimensional scaling showing Bray-Curtis dissimilarity indices over 42 immunologic variables. Variation explained by each axis is indicated in the parentheses. (**C**) All individuals were plotted with multidimensional scaling showing Bray-Curtis dissimilarity indices over 22 immunologic variables, selected for significant variance between groups. Variation explained by each axis is indicated in the parentheses.

JIA is a heterogeneous disease. To determine whether the JIA immune signature identified above showed separation based on disease characteristics, we classified JIA patients into subsets. JIA patients were initially classified along the ILAR guidelines, into oligoarticular persistent JIA (n=15), oligoarticular extended JIA (n=13), RF-polyarticular JIA (n=33), RF+ polyarticular JIA (n=2), enthesitis-related JIA (n=3) and systemic JIA (n=19). The characteristics of the patient cohort as given in **Table 1**. Due to the small number of patients in some ILAR classifications, and the more recent observation that ANA positivity distinguishes a relatively homogeneous group irrespective of the number of affected joints (22), we also used a second classification system, an ANA-based classification. This led to a final grouping of 22 oligoarticular/polyarticular ANA negative (ANA-), 43 oligoarticular/polyarticular ANA+, and 19 systemic JIA patients (**Supplementary Table 3**). Finally, we used a third combined classification strategy, which grouped the patients as persistent oligoarticular (n=15), combined polyarticular and extended oligoarticular (n=48) and systemic JIA (n=19) (**Supplementary Table 4**). As alternative grouping systems exist for JIA, we fully annotated the dataset, allowing independent analysis by alternative criteria (**Supplementary Dataset 1 and 2**). PCA analysis of JIA patients according to ILAR classification showed complete overlapping of most JIA subsets, with increased separation of the RF polyarticular group (limited in number) and the systemic JIA group (**Figure 2A**). Likewise, classification based on ANA status (**Figure 2B**) and the disease-severity classification (**Figure 2C**) also showed almost complete overlap, with only the systemic JIA cluster separating. In each case this systemic cluster showed enhanced separation from the healthy controls along the JIA disease-axis, consistent with sJIA manifesting as a more extreme polarization of the JIA immune signature. As the primary separation was between JIA patients and healthy individuals, we reclustered JIA subsets in the absence of healthy individuals (**Figure 2D-F**). Again, while there was a trend for systemic JIA to separate from the other subtypes, the overall immune deviation signature of JIA was independent of JIA clinical subtype. Together these results suggest that JIA patients, while clinically distinct, share common immunological perturbations.

**Table 1.** Characteristics of JIA patients classified by ILAR criteria. OA, oligoarticular; PA, polyarticular; Can, Canakinumab; Toc, Tocilizumab; ETN, Etanercept; Ada, Adalimumab; Aba, Abatacept.

|  | Healthy controls | Oligoarticular persistent | Oligoarticular extended | RF-polyarticular | RF+ polyarticular | Enthesitis-related | Systemic JIA |
| --- | --- | --- | --- | --- | --- | --- | --- |
| Total N | 43 | 15 | 13 | 33 | 2 | 3 | 19 |
| Male, n (%) | 16 (37.2) | 2 (13.3) | 1 (7.7) | 8 (24.2) | 1 (50) | 1 (33.3) | 7 (36.8) |
| **Age at time of sampling (years)** |  |  |  |  |  |  |  |
| Median (Interquartile range) | 9 (4.75-12) | 9 (4.5-10.5) | 8 (6-11) | 8 (6-10) | 13 (11.5-14.5) | 14 (13-14.5) | 12 (6.5-20.5) |
| Range | 2-17 | 2-16 | 2-13 | 2-18 | 10-16 | 12-15 | 0.75-34 |
| **Years of disease duration** |  |  |  |  |  |  |  |
| Median (Interquartile range) | - | 3.45 (1.7-6.9) | 3.4 (2.3-7) | 2.2 (0.9-5.15) | 3.4 (2.1-4.7) | 4.4 (2.85-5.95) | 6.4 (2-12.05) |
| Range | - | 0.2-13.2 | 0.1-10.7 | 0.3-8.3 | 0.8-6.0 | 1.3-7.5 | 0-21.1 |
| **Disease properties** |  |  |  |  |  |  |  |
| Patients with ANA, n (%) | - | 10 (66.7) | 12 (92.3) | 20 (60.6) | 1 (50) | 0 | 5 (26.3) |
| sJIA patients with MAS, n (%) | - | - | - | - | - | - | 5 (26.3) |
| Patients with active disease, n (%) | - | 8 (53.3) | 10 (76.9) | 13 (39.4) | 2 (100) | 1 (33.3) | 8 (42.1) |
| **JIA medication** |  |  |  |  |  |  |  |
| No medication or nonsteroidal anti-inflammatory drugs, n (%) | - | 7 (46.7) | 3 (23.1) | 9 (27.3) | 0 | 2 (66.7) | 1 (5.3) |
| Methotrexate or leflunomide without biologics, n (%) | - | 6 (40) | 8 (61.5) | 22 (66.7) | 1 (50) | 0 | 5 (26.3) |
| Biologics (Can, Toc, ETN, Ada, Aba ± methotrexate or leflunomide), n (%) | - | 2 (13.3) | 2 (15.4) | 2 (6.1) | 1 (50) | 1 (33.3) | 13 (68.4) |
| ETN and/or Ada, n (%) | - | 2 (13.3) | 1 (7.7) | 2 (6.1) | 1 (50) | 1 (33.3) | 2 (10.5) |
| Can, n (%) | - | 0 | 0 | 0 | 0 | 0 | 3 (15.8) |
| Toc, n (%) | - | 0 | 0 | 0 | 0 | 0 | 8 (42.1) |
| Aba, n (%) | - | 0 | 1 (7.7) | 0 | 0 | 0 | 0 |
| **Dose of steroids (Medrol) (mg/kg)** |  |  |  |  |  |  |  |
| Median (Interquartile range) | - | 0 (0-0) | 0 (0-0) | 0 (0-0.03) | 0 | 0 | 0.05 (0-0.3) |
| Range | - | 0-0.25 | 0-0.3 | 0-0.3 | 0 | 0 | 0-1.4 |

**Figure 2.**
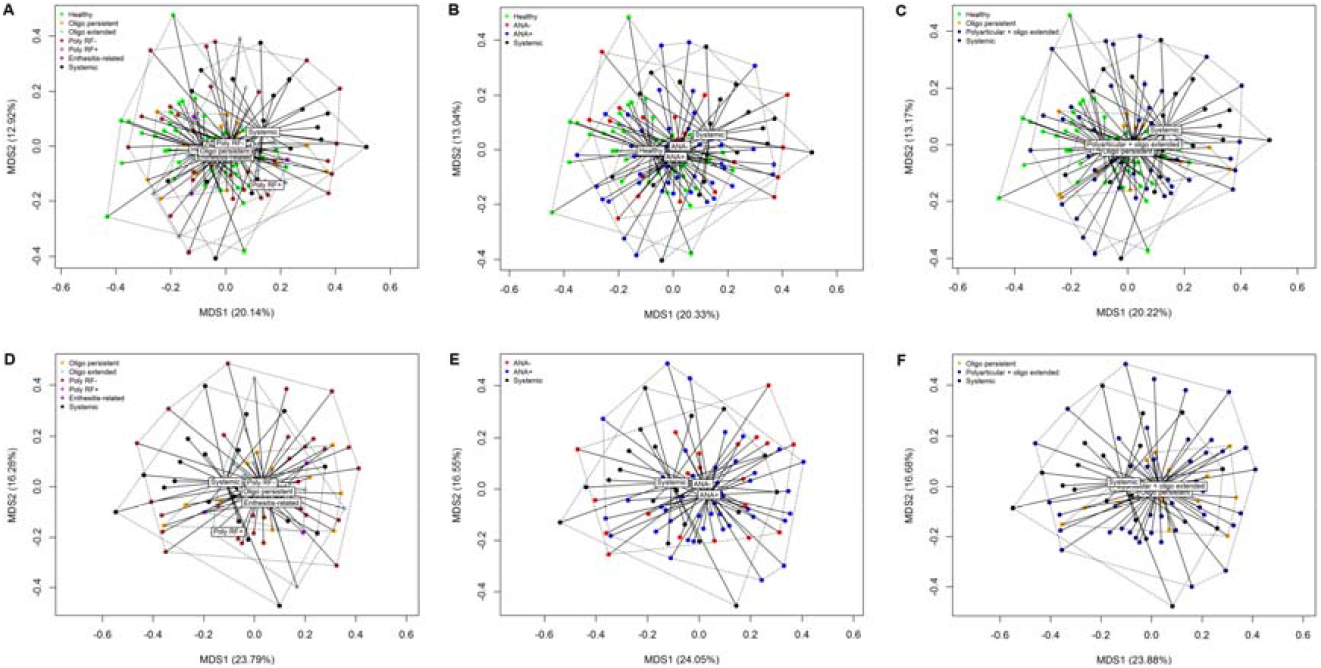
Juvenile Idiopathic Arthritis subsets share a common immunological disturbance. Healthy controls (n=43) and JIA patients (n=85) were assessed for immune phenotype. All individuals were plotted with multidimensional scaling over 42 immunologic variables. (**A**) Healthy individuals clustered with JIA patients, classified upon ILAR criteria. (**B**) Healthy individuals clustered with JIA patients, classified based on ANA status. (**C**) Healthy individuals clustered with JIA patients, classified based on combined classification. (**D**) JIA patients, classified upon ILAR criteria, clustered alone. (**E**) JIA patients, classified upon ANA status, clustered alone. (**F**) JIA patients, classified upon combined classification, clustered alone. Individuals were plotted with multidimensional scaling showing Bray-Curtis dissimilarity indices over 42 immunologic variables. Variation explained by each axis is indicated in the parentheses.

### Juvenile Idiopathic Arthritis patients present with a disturbed adaptive immune system

To unravel the individual drivers of the global immune profile shifts in JIA, we assessed each parameter individually. Using a stringent statistical analysis, and taking into account multiple testing, significant changes were observed in 22 distinct immunological parameters in JIA patients (**Figure 3**). Compared to healthy controls, the disease-control patients registered 17 significant changes in individual immunological parameters (**Figure 3**), notably with 16 of these 17 changes overlapping with the JIA-changed parameters. While there was a large overlap in the changed immunological parameters between JIA and disease controls, the degree of change shifted, with some inflammation-associated changes much stronger in JIA than non-JIA patients, and vice versa. Thus we can consider a shared set of immune phenotypes which are sensitive to inflammatory disease in children (a universal inflammatory signature), while the exact constellation of changes can be disease-specific, driving the separation on a PCA plot (**Figure 1C**).

**Figure 3.**
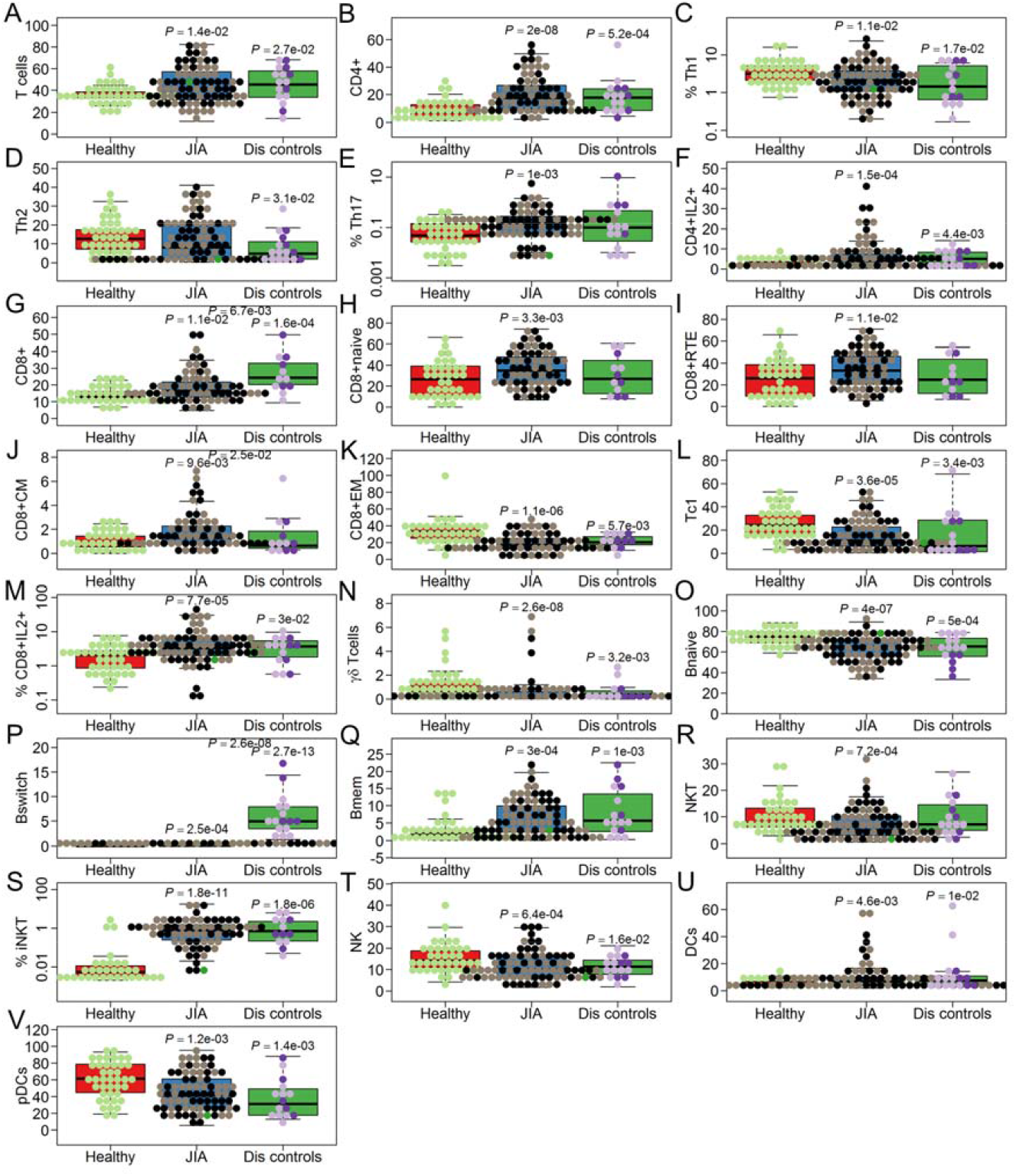
Shared and distinct immunological profiles of JIA and systemic inflammation. Healthy controls (n=43), JIA patients (n=85) and disease controls (n=16) were assessed for immune phenotype. JIA active disease status is indicated by the black dots, quiescent disease is shown by empty dots. Within JIA patients, no disease status was available for one individual indicated by green. Within disease controls, SLE patients are indicated by dark purple and systemic inflammatory disease is indicated by light purple. Box plots for each significant immune parameter are shown, non-significant parameters are not shown. (**A**) T cells, (**B**) CD4+, (**C**) Th1, (**D**) Th2, (**E**) Th17, (**F**) CD4+IL2+, (**G**) CD8+, (**H**) CD8+naïve, (**I**) CD8+RTE, (**J**) CD8+CM, (**K**) CD8+EM, (**L**) Tc1, (**M**) CD8+IL2+, (**N**) yö T cells, (**O**) Bnaive, (**P**) Bswitch, (**Q**) Bmem, (**R**) NKT, (**S**) iNKT, (**T**) NK, (**U**) DCs and (**V**) pDCs. Boxes and center lines represent interquartile range (IQR) and median, respectively; whiskers, 1.5× IQR. Statistical comparison was based on Kruskal-Wallis one-way ANOVA followed by Dunn post-hoc test, adjusted with the False Discovery Rate method. *P* values above patient groups indicate significant difference as compared to healthy controls, *P* values between patient groups indicate significant differences between the JIA and disease controls.

Among the significant changes observed in the immunophenotype of circulating cells in JIA patients, several were suggestive of important biological changes. First, JIA patients displayed increased numbers of CD4 T cells (**Figure 3B**), with particular increases in IL-17 and IL-2 secreting CD4 T cells (**Figure 3E,F**). The increase of Th17 cells, in conjunction with prior evidence (6-8), indicates that activation of CD4 T cells contributes to the inflammatory manifestations. Changes in the CD8 compartment, by contrast, were more consistent with a decreased, rather than enhanced, CD8 effector response. CD8 T cells were raised in JIA patients (**Figure 3G**), however there was a significant increase in naïve and recent thymic emigrant CD8 T cells (**Figure 3H,I**). Notably, these increases were unique to JIA, and not observed in the disease control cohort. Within antigen-experienced CD8 T cells, there was an increase in central memory (Figure 3J) and IL-2-producing CD8 T cells (**Figure 3M**) at the expense of IFNγ producing (Tc1) or effector memory CD8 T cells (**Figure 3K,L**). The decrease in IFNγ-producing CD8 T cells is of particular note, with the standard model of progression of sJIA into Macrophage Activation Syndrome driven by excessive production of IFNγ by effector memory CD8 T cells (23). Thus, while IFNγ has been proposed as a key pro-inflammatory cytokine in JIA, our data in the peripheral blood is more consistent with impeded production of IFNγ. Altered activation of T cells in JIA patients was accompanied by activation of B cells, with a decrease in naïve B cells (**Figure 3O**), coupled with an increase in switched B cells (**Figure 3P**) and memory B cells (**Figure 3Q**). The degree of B cell activation was, however, much lower in JIA than disease controls, where a subset of patients exhibited a profound increase (**Figure 3P**). Beyond these major adaptive cell types, JIA patients displayed a relative decrease in γδ T cells, NK cells, plasmacytoid dendritic cells, while iNKT cells were increased (**Figure 3**). Together, these results suggest a pathophysiological process in the circulating lymphocytes involving the suppression of IFNγ production by CD8 T cells and excessive CD4 T cell differentiation into the Th17 lineage.

When JIA subsets were considered, the changes observed at the individual parameter level reflected the analysis at the global level: the immunological changes occurring in JIA versus healthy children were highly similar among JIA subtypes. In general, immunological changes in each JIA subset mirrored the others in trend, if not statistical significance, regardless of whether the ILAR criteria (**Supplementary Figure 2**), ANA-based criteria (**Supplementary Figure 3**) or combined criteria (**Supplementary Figure 4**) were used. An exception was found with sJIA, which showed a more profound increase in CD4+ T cells and in naïve CD8 T cells than the other JIA subsets (**Supplementary Figures 2–4**), and the cluster of ANA+ polyarticular and oligoarticular JIA patients, which were the only to show increased switched B cells. Together, these results indicate that, despite the different clinical manifestations, JIA subtypes share a distinct immunophenotype, with only relatively minor immunological changes correlating with the particular JIA clinical subtype.

Potential confounding factors in the immunological analysis of JIA patients include disease activity and immunological changes secondary to treatment. As JIA subtypes manifested a mostly similar immune phenotype, or at least phenotypic changes along the same spectrum, we merged JIA subtypes in an attempt to control for disease activity. Separation of JIA patients into patients that had either active disease or were quiescent revealed a more extreme immune signature in JIA patients that were sampled at a time point of active disease, although even JIA patients sampled during disease inactivity showed separation from the healthy controls (**Figure 4A**). The immunological relationships observed in patients with inactive JIA were largely replicated in patients with active JIA (**Figure 4B**), however active JIA patients did show altered interrelatedness for several immune parameters. For example, in inactive JIA patients, no relationship was observed between CD8 T cells and memory B cells, however a strong positive relationship emerged in active JIA patients (**Figure 4B**). At the individual parameter level, of the 21 immune parameters that showed a significant difference between JIA and healthy, 20 showed no change between patients sampled at active or inactive disease stages (**Supplementary Figure 5**). Only one parameter, NK cell percentage, showed a significant difference based on disease state, with the drop in NK frequency only observed in the patients with inactive disease (**Figure 4C**). Our findings suggest that disease activity correlates with quantitative rather than qualitative changes in the signature immune dysregulation, with the underlying JIA immune dysfunction remaining present in patients with disease control.

**Figure 4.**
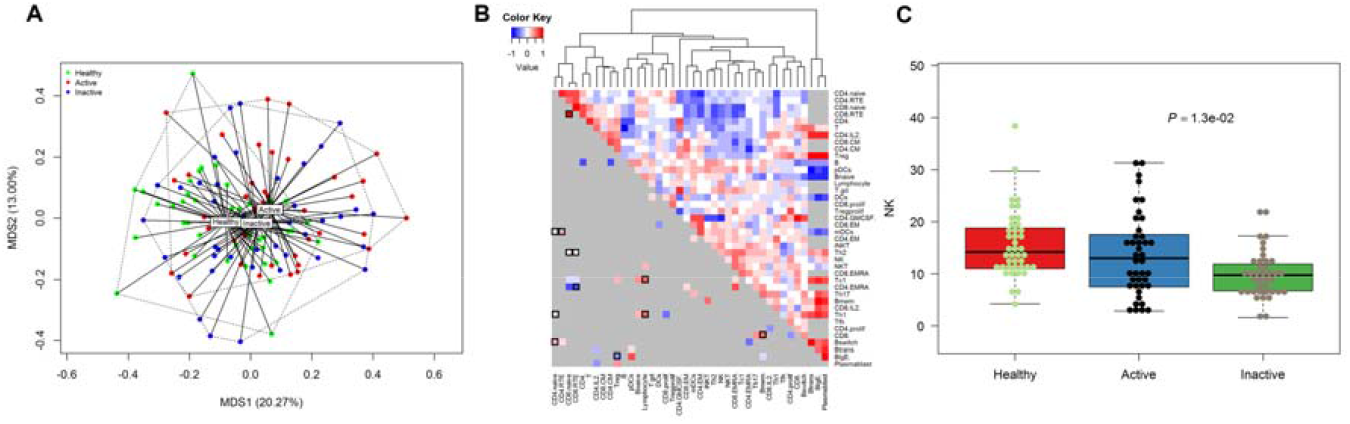
Immunological architecture of JIA is maintained into disease remission. Healthy controls and JIA patients were assessed for immune phenotype. (**A**) Healthy controls (C; n=43), JIA patients with quiescent disease at sampling (inactive; n=43) and JIA patients with active disease (JIA active; n=42 plotted with multidimensional scaling over 42 immunologic variables. All individuals were plotted with multidimensional scaling showing Bray-Curtis dissimilarity indices over 42 immunologic variables. Variation explained by each axis is indicated in the parentheses. (**B**) Upper right, above the diagonal: coregulation between pairs of cell types in JIA patients with inactive disease (n=42) (red: positive correlation coefficient, blue: negative correlation coefficient, light gray: no data available). Unbiased clustering of coefficients was performed to group coregulated cell types. Lower left, below the diagonal: dark gray indicates coregulation between pairs of cell types in JIA patients with active disease (n=42 that is preserved from inactive JIA patients). Coregulation between pairs of cell types that are significantly altered by disease activity (p < 0.05, and boxed if *p* < 0.01) are colored (red: positive correlation coefficient, blue: negative correlation coefficient). (**C**) Box plots NK cells shown. Boxes and center lines represent interquartile range (IQR) and median, respectively; whiskers, 1.5× IQR. Statistical comparison was based on Kruskal-Wallis one-way ANOVA followed by Dunn post-hoc test, adjusted with the False Discovery Rate method.

To assess the contribution of medication to the immune phenotype, we grouped JIA patients into those who were without medication (or on non-steroidal anti-inflammatory drugs only), those who were being treated with methotrexate or leflunomide, and those who were on biologic therapies (including TNF blockade, anti-IL1, anti-IL6 and abatacept, with or without additional non-biologic therapy). Using a global analysis, the greatest separation was observed between healthy controls and patients on either steroidal or biological treatment, with untreated patients showing intermediate clustering (**Figure 5A**). This effect was reflected at the individual parameter level, where more pronounced changes were observed in patients undergoing steroidal or biologic treatment (**Figure 5B**). The same result was observed when treatment was considered for each JIA subtype, albeit with reduced statistical power (**Supplementary Figure 6**). While this effect paralleled that of disease activity, the effects were independent: no segregation of disease active / inactive patients was observed within the treatment categories (**Figure 5B**). The effect could be partially explained, however, by the discrete clustering of sJIA patients, which, regardless of disease activity, showed a more extreme immune signature, and which were heavily enriched for patients on steroidal or biologic treatment. Thus, when only non-sJIA patients were classified by treatment, a lower degree of separation between treatment groups was observed (**Figure 5C**). Together, this data supports the contention that the immune phenotype observed above is primarily associated with JIA diagnosis rather than (secondary to) treatment.

**Figure 5.**
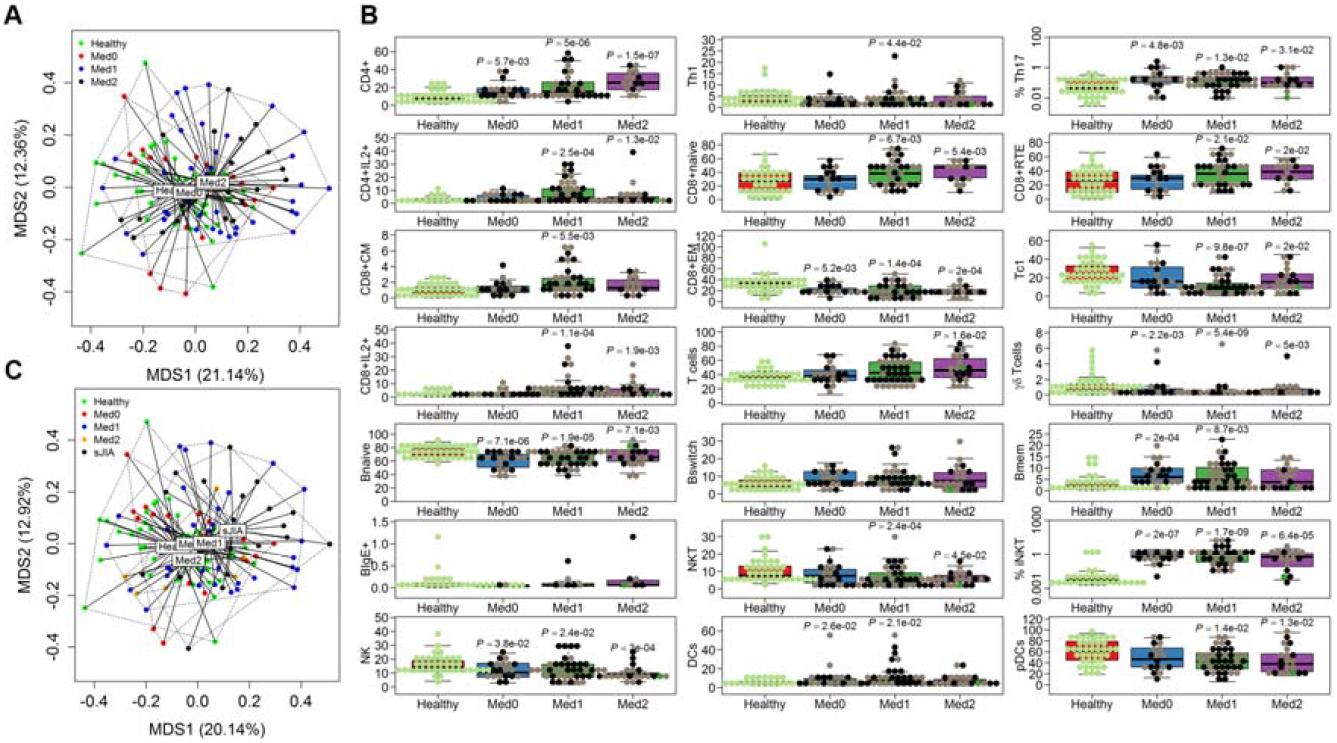
Immunological architecture of JIA is not primarily driven by medication. Healthy controls and JIA patients were assessed for immune phenotype. (**A**) Healthy controls (C; n=43), untreated JIA patients (Med0, no medication or nonsteroidal anti-inflammatory drugs; n=22), steroid-treated JIA patients (Med1, methotrexate or leflunomide; n=42), or biologic-treated JIA patients (Med2, Canakinumab, Tocilizumab, Etanercept, Adalimumab, Abatacept, with or without additional methotrexate or leflunomide treatment; n=21) were plotted with multidimensional scaling over 42 immunologic variables. All individuals were plotted with multidimensional scaling showing Bray-Curtis dissimilarity indices over 42 immunologic variables. Variation explained by each axis is indicated in the parentheses. (**B**) Immune phenotypes in which a significant change was observed in one or more JIA subtype were assessed for correlates with treatment status. Active disease status is indicated by the black dots. Boxes and center lines represent interquartile range (IQR) and median, respectively; whiskers, 1.5× IQR. P values show significant difference as compared to the controls. Statistical comparison was based on Kruskal-Wallis one-way ANOVA followed by Dunn post-hoc test implemented in R and *P* values were adjusted with the False Discovery Rate method.

### Immune-aided machine learning can discriminate between JIA patients and healthy children

A key application of immunophenotyping in JIA would be to drive assay-based personalized therapeutic choice of treatment strategy. While such an outcome requires a multi-year longitudinal immunophenotyping dataset, we sought to determine in the current dataset whether machine learning could identify the primary immunological features of JIA, as a basis for future discovery. Three approaches were trialed: random forests, artificial neural networks and support vector machines; with the random forest approach showing superior performance in separating healthy individuals from JIA patients (**Supplementary Table 5**). A random forest is an ensemble of deep decision trees, where each tree is constructed using a random subset of samples. The trees are constructed such that every split only considers a random subset of features to avoid the selection of only closely correlated features and allow the consideration of alternative features (12, 13). The data was split into 10 random folds, with hyperparameter training and selection on 9 folds and testing on the 10^th^, with rotation of the testing fold. After the construction of 10,001 trees, the optimal random forest selection strategy was capable of discriminating JIA patients from healthy controls with an area under the curve of 89.6% (**Figure 6A**). The key contributing feature to JIA “diagnosis” was the increased number of iNKT cells (**Figure 6B**). Indeed, the increased frequency of iNKT cells alone resulted in an area under the curve of 91.2% (**Figure 6A**), superior to the random forest in accuracy, although reliant on a single parameter. This result demonstrates the utility of machine learning in prioritizing identified changes for explanatory power, beyond *a priori* biological rationale, with potential for use in the design of diagnostic assays.

**Figure 6.**
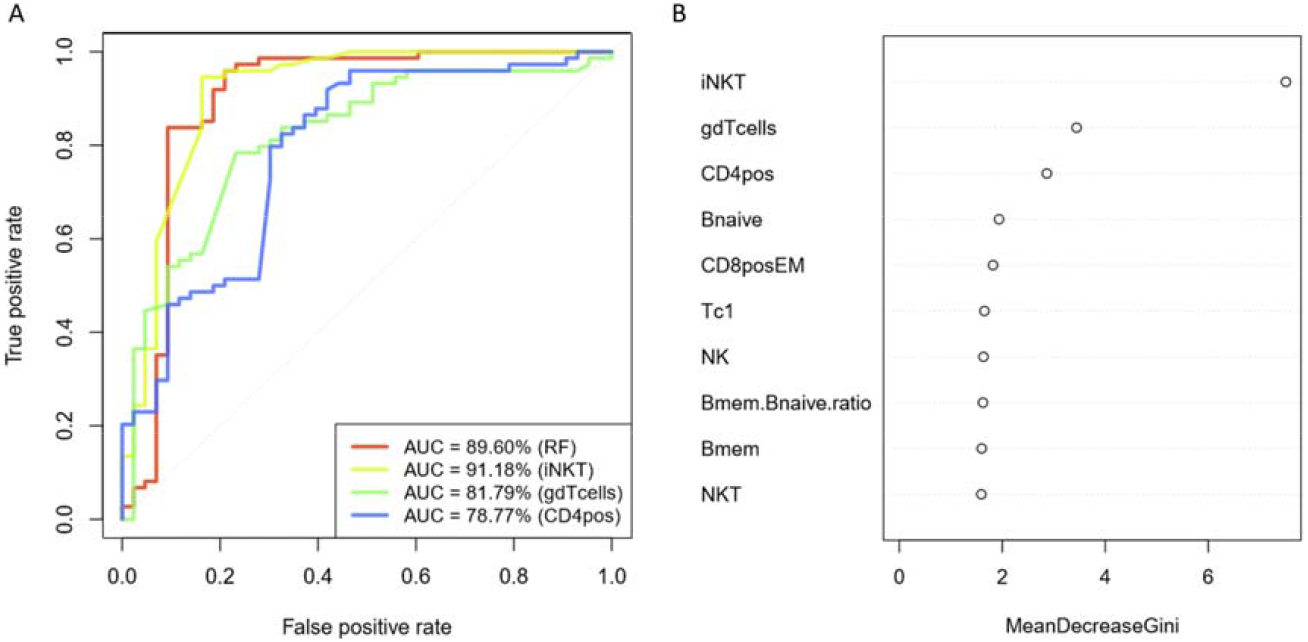
Machine learning identifies iNKT cells as primary predictive immunological change in JIA. Random forests were trained on immunological data from healthy controls (n=43) and JIA patients (n=72), selecting for capacity to discriminate between the two groups. (**A**) ROC curves computed from the out-of-bag predictions of a random forests trained on the entire dataset (red curve) and the top three features identified by the trained random forests (other curves). (**B**) The top ten features in the random forest trained on the entire data set, and contribution to the optimal random forest approach.

## Discussion

Improved understanding of the immunological architecture of JIA is required for pathophysiology-based diagnosis and treatment. Through extensive immune phenotyping of the adaptive immune system, we found an immunological signature of JIA in a large cohort of Belgian children. This signature comprised two components: first, a shared “universal signature” of inflammation, common between children with JIA, SLE or diverse inflammatory diseases. Such a universal signature is unlikely to derive from shared pathophysiology, and may instead reflect a signature response of children’s immune systems to inflammatory disease. Second, a set of JIA-specific immune changes drove a deviance from the universal signature, providing an axis of discrimination between JIA and the non-JIA inflammatory diseases tested here. While the distinct clinical presentation of these diseases does not necessitate biologically-derived discrimination, such changes may identify the disease-specific pathophysiological processes.

Within the JIA cases here assessed, the immunological signature was largely shared across disease subtypes. The JIA signature was enhanced in two populations: JIA patients sampled at a time-point of active disease, and JIA patients of the systemic subtype. sJIA has long been considered an entity distinct from other, more common, JIA subtypes, mediated by abnormalities in the innate immune system with features of an autoinflammatory disease. Indeed, high-depth immunophenotyping of the innate immune system recently identified a sJIA signature response to stimuli (10). This simple innate versus adaptive division between sJIA and other JIA subtypes is, however, challenged by the genetics, with association of the HLA-DRB*11 class II allele to sJIA susceptibility indicating a strong role for the adaptive immune system (24). How then to reconcile these datasets? A biphasic clinical course of sJIA was recently proposed where innate immune mechanisms dominate at disease onset, eliciting systemic inflammation through increased levels of IL-6, IL-18, S100A8/A9, S100A12 and IL-1ß, followed by a second phase where the adaptive immune system mediates chronic arthritis (25). Integrating our data with prior studies, the innate-drivers at disease onset sJIA-specific (10), while the ongoing adaptive disturbances may reflect process in line with, but exaggerated from, non-systemic JIA patients.

The pathophysiological process of JIA disease susceptibility identified through our analysis is one of complex immune network failure, rather than the generation of a single pathology-driving event. A key interaction may be the balance between Th1/Tc1 and Th17 cells. Th17 cells are pathogenic in mouse models, with IL-17- and IL-23-deficiency invoking resistance to arthritis induction (26-29). Evidence from human studies over the past 20 years provides ample data to support Th17 cells as drivers of autoimmunity in JIA and RA. Synovial fluid of both JIA and RA patients was shown enriched for Th17 cells (30-33) and elevated plasma levels of IL-17 in oligo- and poly-JIA were shown to correlate with disease activity (34). Addition of IL-17 to human *ex vivo* models enhanced IL-6 production and collagen destruction, while inhibiting collagen synthesis by RA synovium explants (35). In contrast to IL-17, the functional role of IFNγ is decidedly ambivalent. While Th1 and Tc1 cells are considered inflammatory, IFNγ also suppresses the differentiation of Th17 cells, which can drive a net suppressive impact on autoimmunity. Indeed, in mouse models, treatment with IFNγ suppresses arthritis development due to impeded Th17 differentiation (36). Conversely, deficiency in IFNγ promotes a sJIA-like disease upon Freund’s complete adjuvant in mice (9), and IL-12 knockout mice, deficient in Th1 cells, are likewise susceptible to arthritis induction (26, 27, 37), both phenotypes attributed to increased production of Th17 cells. A primary defect in IFNγ-production could thus drive a pro-Th17 state that is permissive for JIA development. Alternatively, the causality could be reversed; IL-17 production reduces Th1 differentiation in RA (38), and thus elevated Th17 numbers may be the primary effect. The identification of strong changes in the NKT cell compartment was the most surprising change observed. As these cells can have pro- or anti-arthritic properties in mice, depending on the model (39), either a potential mechanistic involvement or a compensatory biomarker function could be envisioned in JIA.

Beyond the mechanistic insights offered into disease pathogenesis, the utility of machine learning to extract JIA diagnosis from the immunological signature serves as a proof-of-principle of immune-directed machine learning in precise diagnosis and personalized therapeutic choice. The dataset used here lies at the threshold of manual and automated analysis, while future studies of larger populations and more extensive immune profiling will require automated analysis for full value extraction. Here a machine-learning approach was able to build a diagnostic algorithm for JIA diagnosis, and, critically, we were able to extract the informative parameters from the diagnostic algorithm. In principle, these limited parameters could be used to design a simplified immunophenotyping assay for JIA diagnosis. In practice, such an assay would be of relatively limited use: JIA diagnosis in secondary and tertiary reference centers is efficient (although diagnosis during primary care may be delayed due to overlapping early symptoms and the lack of specific biomarkers (40), and the machine-learning algorithm was derived from comparison to healthy individuals rather than patients with a relevant confounding disease (such as infectious arthritis). None the less, the ability of machine learning to extract the relevant signature validates this approach for larger longitudinal cohorts, with deeper immunophenotyping, with a focus on personalized response to treatment. The features of unbiased analysis and identification of features with combinatorial diagnostic power would allowed an immune-led machine learning process to identify those immune signatures predictive of efficient response to particular treatments. Such an immune-driven personalized medicine approach to JIA would greatly improve the appropriate treatment selection for patients.

## Acknowledgments

This work was supported by the ERC Start Grant IMMUNO. E.V.N., V.L., L.V.E. and S.H-B. were supported by fellowships from the FWO. The authors thank Isabelle Meyts for the provision of disease control samples.

